## Supplementary file for "DYRK1A Interacts with the Tuberous Sclerosis Complex and Promotes mTORC1 Activity"

**^3^** National Facility for Protein Science in Shanghai, Zhangjiang Lab, Shanghai, 201210, China.

**^4^** Stowers Institute for Medical Research, 1000 East 50th Street, Kansas City, Missouri 64110, USA.

**^5^** Department of Cancer Biology, The University of Kansas Medical Center, 3901 Rainbow Boulevard, Kansas City, Kansas 66160, USA.

**^6^** CSIR–Centre for Cellular and Molecular Biology, Habsiguda, Uppal Road
Hyderabad, 500007, India.

**^7^** Department of Biochemistry and Molecular Cell Biology, Shanghai Key Laboratory of Tumor Microenvironment and Inflammation, Shanghai Jiaotong University School of Medicine, Shanghai 200025, China.

† These authors contributed equally to this work

‡ Current address: Centre for Writing and Pedagogy, School of Interwoven Arts and Sciences, Krea University, Sri City, AP, 517647, India

**Keywords:** Microcephaly, Cell growth, Minibrain, Drosophila, Neuromuscular Junction
**Figure S1.** Analysis of cell size after induction of DYRK1A expression with increasing dosage of Doxycycline.

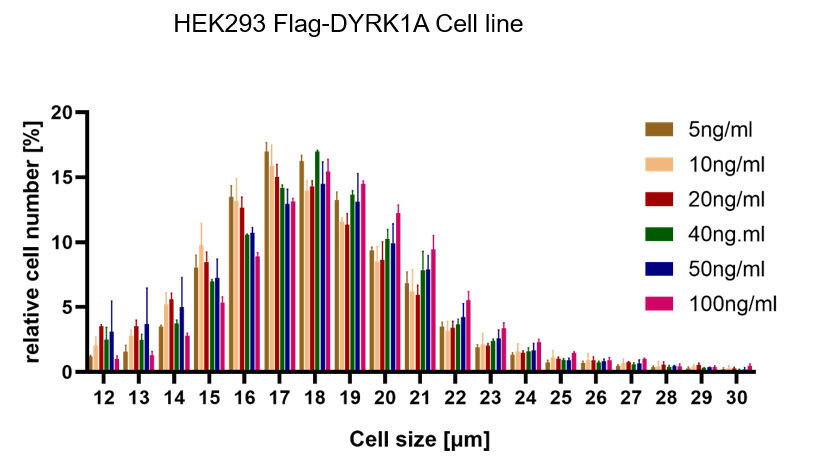

**Figure. S2.** TSC1/TSC2 interact with DYRK1A. (A) Flag and HA beads were used to pull-down HA3-TSC1 and Flag-TSC2 from whole cell extracts of HEK293 transfected with HA3-TSC1 and Flag-TSC2. Blots were probed with Flag, HA, and DYRK1A antibodies. Actin was used as loading control.

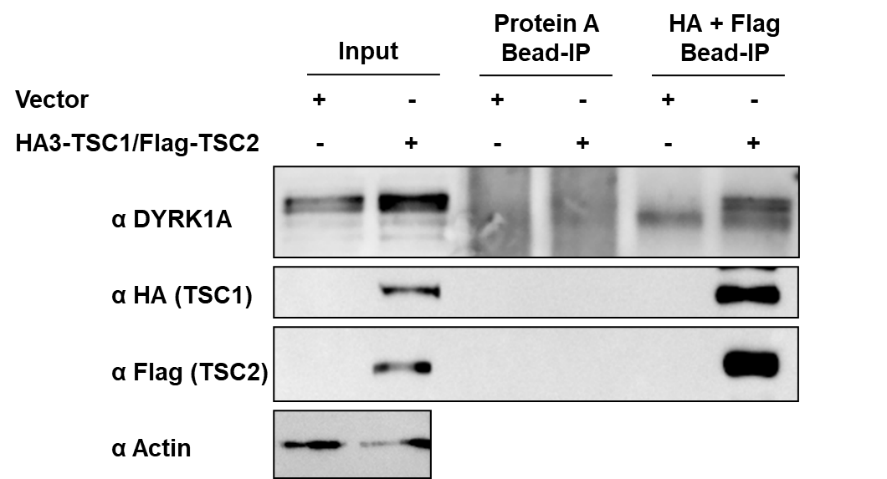

**Figure. S3.** DYRK1A kinase domain interacts with TSC1. (A) Schematic of DYRK1A truncated constructs. All the truncated DYRK1A forms carry Flag tag at the N-terminus. (B) Flag-DYRK1A and Flag-DYRK1A truncated constructs were affinity purified using Flag-beads from HEK293 cells co-transfected with HA3-TSC1 and probed with α-HA and α-Flag antibodies. All DYRK1A constructs, except with deletions in the kinase domain, immunoprecipitated HA3-TSC1. Note that Flag-DYRK1A kinase domain deletion constructs were expressed at lower levels than other constructs. Actin was used as lysate control.

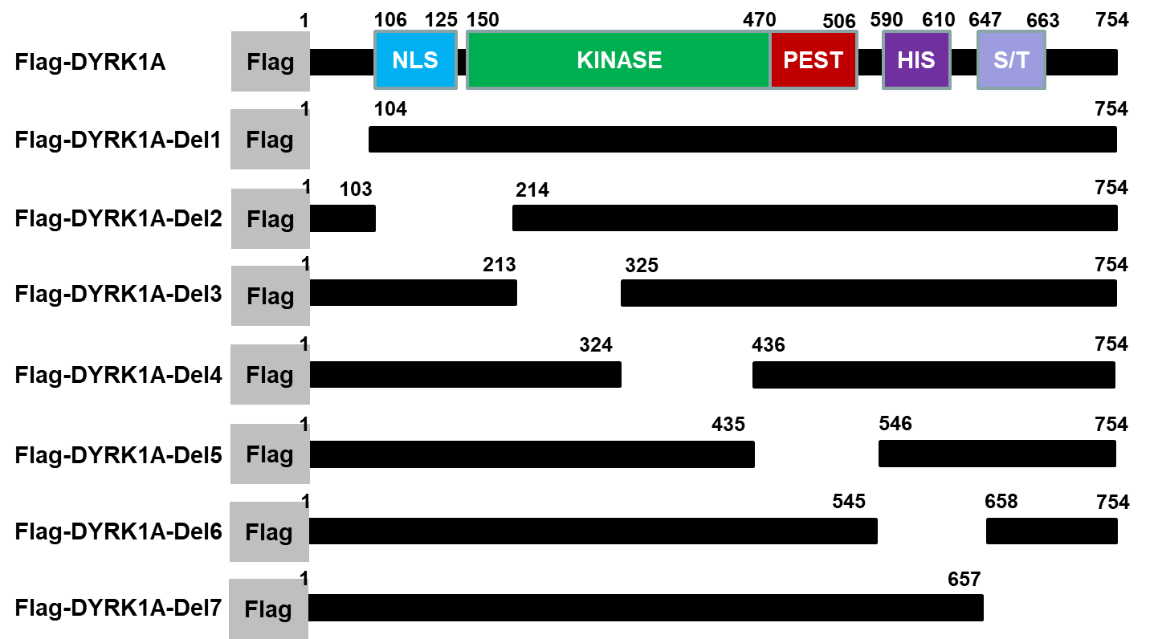

**A**

**B**

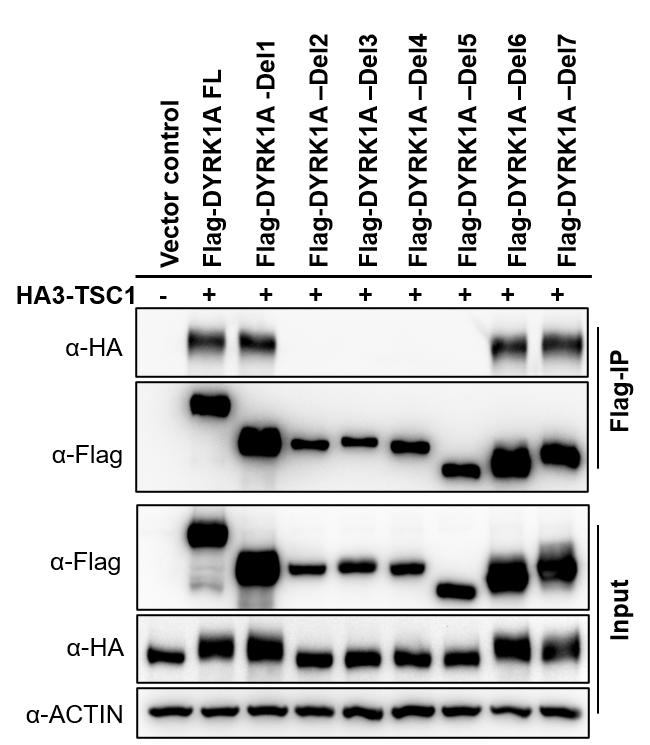

**Fig. S4. mTORC1 inhibitors block the increase in cell size mediated by DYRK1A.**

HEK293 cells expressing *Flag-DYRK1A* and the parental cells were treated with 40ng/ml Doxycycline. At 24 hours mTOR inhibitors Torin1/Rapamycin were added and the cells were further incubated for 24 hours. Data represent the mean ± SD (n = 3 biological replicates).

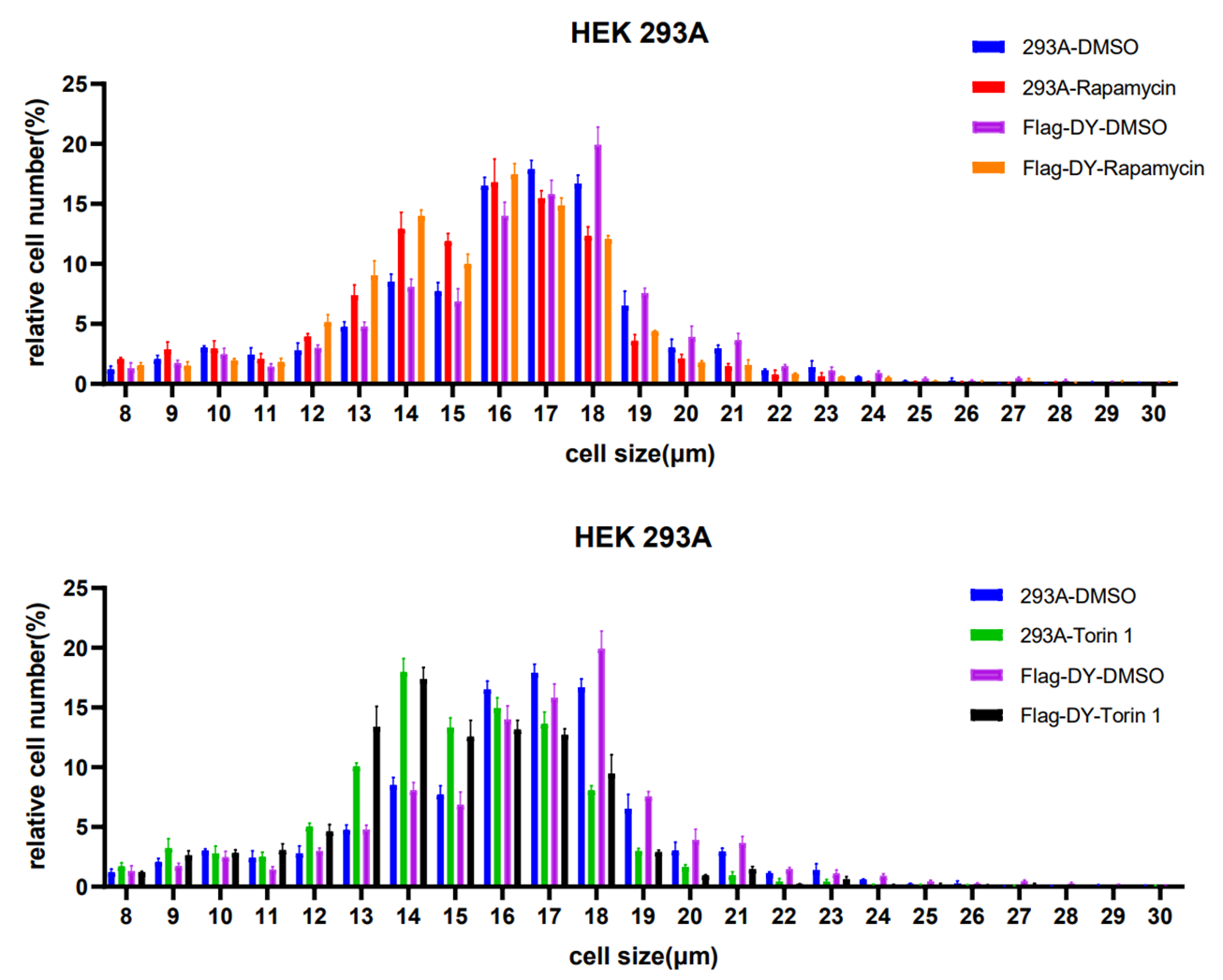

**Fig. S5.** NMJ phenotypes due to TOR gain or loss. NMJ (muscles 6/7) are stained using anti-HRP (Green) and anti-Dlg (Red). Muscles are stained with phalloidin (Blue, A-F). HRP (green) stains the entire neuron and Dlg (red) stains only boutons (Red+Green). (G-I, M) Quantification of bouton numbers - normalized to muscle area (Bouton-NMA). Error bars represent standard deviation. Statistical significance *(p*-values: *** < 0.001; ** < 0.01; * < 0.05) is calculated by unpaired student's *t-*test. (A-C) *gig^109^* alleles show increased bouton numbers (B) as compared to wild-type (WT, Canton S) control (B). Data is quantified in C. (D-F) Expression of dominant negative TOR (*D42-Gal4*>UAS-TOR.ted*,* E) decreases bouton numbers as compared to control (*D42-Gal4*/+, D). D42-Gal4 is a motor neuron specific driver. Data is quantified in F.

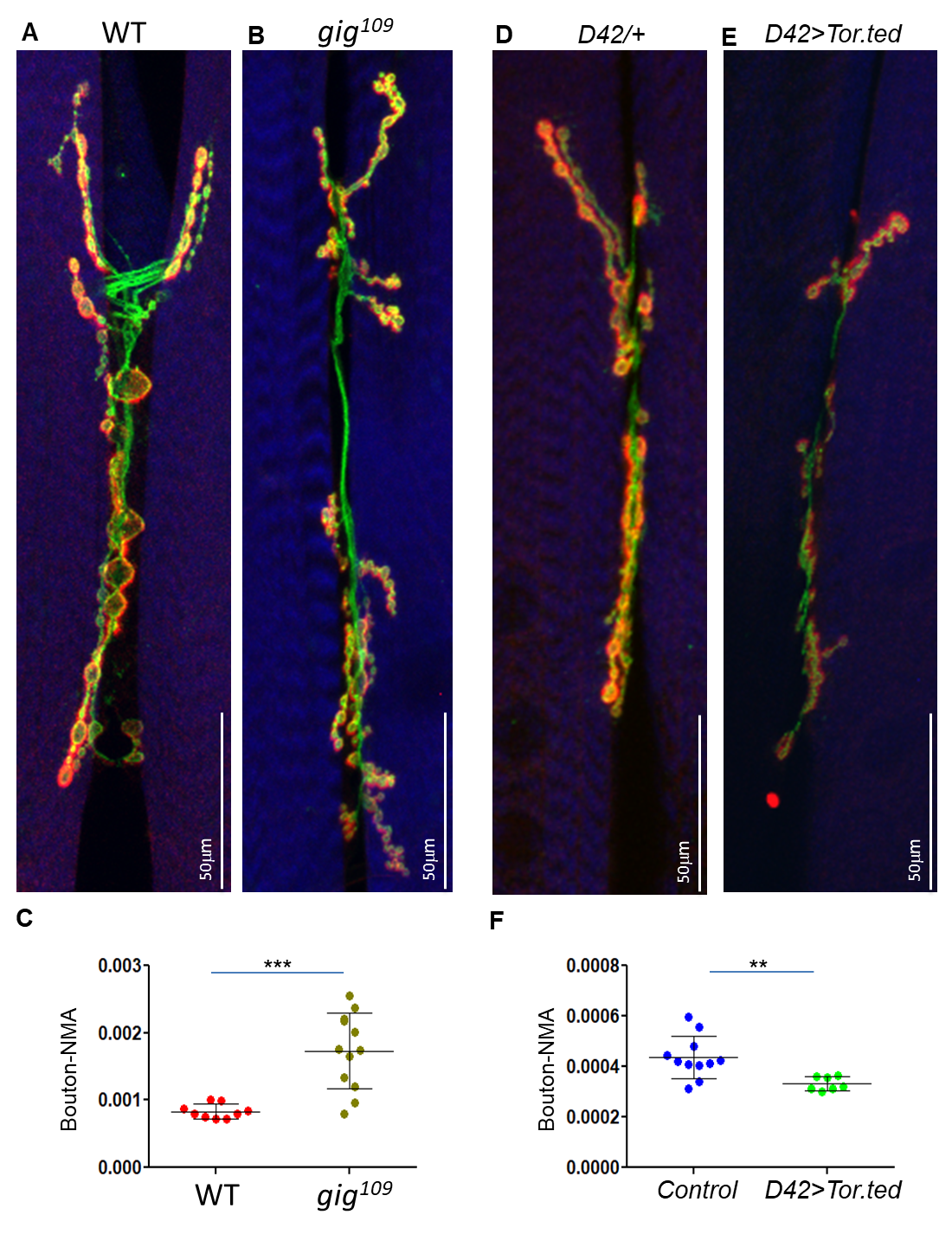

**Table S1.** SgRNA target sequences for mouse cells

| Gene name | SgRNA Target |
| --- | --- |
| Control-sgRNA | Gcgaggtattcggctccgcg |
| Dyrk1a-sgRNA1 | gcgcttttatcggtctccag |
